# Evolutionary analysis of mammalian Rem2, a member of the RGK (Rem, Rem2, Rad, and Gem/Kir) family of small GTPases, reveals the role of selection and epistasis in shaping protein functional constraints

**DOI:** 10.1101/2023.08.28.555117

**Authors:** Alexander G Lucaci, William E Brew, Sergei L Kosakovsky Pond, Anna R Moore

## Abstract

*Rad And Gem-Like GTP-Binding Protein 2 (Rem2)*, is a member of the RGK family of Ras-like GTPases and has been identified in various mammalian species. *Rem2* has been implicated in Huntington’s disease and Long QT Syndrome and is highly expressed in the brain and in endocrine cells. In this study, we examined the evolutionary history of *Rem2* across mammals, focusing on the role of purifying selection and epistasis in shaping its sequence and structure. In our analysis of *Rem2* sequences across 175 mammalian species, we found evidence for strong purifying selection in 70% of non-invariant codon sites of the protein, characteristic of essential proteins that play critical roles in biological processes and is consistent with *Rem2*’s role in the regulation of neuronal development and function. We inferred epistatic effects in 49 pairs of coevolving codon sites in *Rem2,* some of which are predicted to have deleterious effects on human health. Additionally, we reconstructed the ancestral evolutionary history of mammalian *Rem2* using protein structure prediction of extinct and extant sequences. This analysis revealed the dynamics of how substitutions that change the genetic distance of Rem2 can impact protein structure in variable regions while maintaining core functional mechanisms. By understanding the selective pressures, protein- and genetic-interactions that have shaped the sequence and structure of the Rem2 protein, we may gain a stronger understanding of its biological and functional constraints.

## Introduction

Brain development is a highly orchestrated sequence of events involving a multitude of molecular pathways. Disruption to these systematic processes can lead to impairments in brain function and disease states, as seen with neurological disorders such as autism, Alzheimer’s disease, Huntington’s disease, and Parkinson’s disease [Kwakowsky et al., 2023]. Regulatory proteins are of particular interest for understanding proper brain function and development. Recent discoveries of key genes associated with neurological diseases include those associated with synaptogenesis, neuronal architecture, and neuronal function [Kwakowsky et al., 2023; Allen et al., 2016]. Rem2 was identified as a key activity-regulated gene important for synapse development and function [Ghiretti et al., 2011, 2014; Moore et al., 2013, 2018], dendritic complexity [Ghiretti et al., 2014, 2011], and intrinsic plasticity [Moore et al., 2018]. Taken together, understanding the evolutionary history of Rem2, in combination with its cellular function, is critical for identifying potential therapeutic targets in disease states.

### An overview of Rem2 gene biology

Rad And Gem-Like GTP-Binding Protein 2 (Rem2), encodes a small Ras-like GTPase within the RGK family which includes Rem, Rad, Gem and Kir proteins [Finlin et al., 2000]. Since its initial discovery as a potent inhibitor of calcium channel activity [Béguin et al., 2001; Correll et al., 2008, Flynn et al., 2010], Rem2 has been identified as one of the key RGK members expressed in the brain. Rem2 is expressed within regions of the hippocampus and basal ganglia [Liput et al., 2016] and functions as an activity-dependent [Flynn et al., 2010] regulator of synapse formation, dendritic complexity, and intrinsic excitability [Ghiretti et al., 2011; Moore et al., 2013, Ghiretti et al., 2014; Moore et al., 2018]. Additionally, at the cellular level, regulatory interactions have been reported between Rem2 and calmodulin [Béguin et al., 2005], L-type and N-type voltage gated calcium channels (VGCC) [Correll et al., 2008, Flynn et al., 2008], and calmodulin-dependent protein kinase II (CaMKII) [Ghiretti et al., 2013, Royer et al., 2018]. Rem2 also modulates the activity of other small GTPases such as Rac1 and RhoA [Gauthier-Rouvière, 1998], which are themselves important regulators of the actin cytoskeleton and neuronal morphogenesis.

Rem2’s involvement with key cellular pathways for neuronal function, its regulation on neuronal architecture, excitability, and high expression in the hippocampus, a brain region known to be associated with memory and learning [Meck et al., 2013], implicate *Rem2* as a target gene of interest for understanding neuronal health and disease states. Recently, mutations in Rem2 were identified in patients with Huntington’s Disease [Nahalka, 2019] and Long QT Syndrome contributing to channelopathies [Chai et al., 2018]. Another member of the RGK family, Gem, which displays similar roles in neuronal function as Rem2 has been implicated in Timothy Syndrome [Krey et al., 2013; Boczek et al., 2015]. Thus, a further understanding of *Rem2* gene evolution and function will provide deeper insight into the cellular role of the RGK subfamily and calcium channel interactions.

### The conservation of Rem2 throughout its evolutionary history

Selection analyses form a common approach in the field of molecular evolution, used to assess the evolutionary forces that have influenced the dynamics of proteins. These analyses primarily involve the estimation of the omega parameter (ω), which is the ratio of non-synonymous (β) to synonymous (α) substitution rates, expressed as ω = β/α [Muse and Gaut, 1994]. Non-synonymous changes can often exert significant influences on the structural configuration and functional operation of a protein. Conversely, synonymous changes do not modify the amino acid at a specific site, yet they can still have subtle effects on fitness through codon usage bias, translation efficiency, or mRNA structural stability. Typically, the rate of synonymous changes is assumed to be the rate of neutral selection acting upon coding sequences and serves as a baseline against which non-synonymous evolutionary rates can be compared. The estimates of ω conveniently quantify the selective pressure acting upon protein-coding genes and can be readily interpreted to discern the type of selective modality at play (e.g., positive diversifying selection with ω > 1, negative selection with ω < 1, or neutral evolution when ω is not significantly different from 1).

Purifying selection acts to remove multiple types of variation from a population, including deleterious mutations. This process is important in maintaining the functional integrity of genes and preventing the accumulation of mutations that could lead to loss of function or disease. In the case of *Rem2,* this may be related to its role in neuronal development and function, such as GTP and calmodulin binding which have shown to be essential for regulating dendritic complexity [Ghiretti et al., 2011; Correll et al., 2008]. Epistasis, i.e., interaction between two or more sites in a gene, may impact the evolution of a gene and play a role in constricting its functions. In the case of *Rem2*, epistasis could play a role in shaping the genetic interactions that regulate its function through its functional domains such as RGK.

### Purpose of this study

Understanding the history of *Rem2* would offer insights into the processes that have shaped this gene and its function, including its relationship with its functional partners in biological networks. Our study uses a comparative genomic approach with the molecular sequences of *Rem2* in different species to identify patterns of conservation and divergence. An evolutionary study of *Rem2* may shed light on the broader properties and dynamics of the RGK family of small GTPases and their roles in regulating cellular functions, as Rem2 and Gem are the only two RGK family members expressed in the brain [Correll et al., 2008].

Exploring potential ancestral functions of Rem2 may illuminate our current understanding of its role in regulating neuronal development and function. Additionally, the evolutionary trajectory of *Rem2* may be influenced not only by the gene itself but also by its interactions with other genes. Therefore, the results from this work could be applied to future studies to aid in the understanding of the molecular mechanisms underlying the genetic networks of neurological disorders associated with Rem2 mutations and to identify potential targets for therapeutic intervention.

## Methods

### Data retrieval

In this study, we queried the NCBI database for the *Rem2* gene using the following URL, https://www.ncbi.nlm.nih.gov/gene/161253/ortholog/?scope=1338369&term=Rem2 (last accessed December 2022). As our primary interest in this study is to explore the evolutionary history of *Rem2* in *Mammalia*, we limited our search to only include species from this taxonomic group. This query returned 186 full gene sets (transcripts and protein sequences) with one sequence per species. We downloaded all available files: Reference sequence (RefSeq) protein sequences, RefSeq transcript sequences, and Tabular data (CSV) which contains sequence metadata. In Table S1, we provide information of the species included in this analysis.

### Data curation

We used the full protein sequence and full gene transcript files (which include 5’ and 3’ UTR) to retrieve coding sequences (CDS). Our process also removed low-quality protein sequences (9 sequences) from analysis as these sequences may bias or contaminate our results and analyses. For example, these sequences had invalid features such as incompletely resolved codons which result in unresolved amino acids and were subsequently excluded from our analysis.

### Multiple sequence alignment

We generated codon-aware alignments for the filtered set of protein sequences by following the procedure available at the codon-msa GitHub repository (github.com/veg/hyphy-analyses/tree/master/codon-msa). We extracted and relied on the Human *Rem2* coding sequence (see Table S1 for accession) as the reference sequence for a reference-based alignment approach. In-frame nucleotide sequences were translated then the protein sequences were aligned with Multiple Alignment using Fast Fourier Transform (MAFFT) v7.505 [Katoh & Standley, 2013], and then were mapped back to their respective codons.

### Filtering outlier sequences

In order to raise our confidence in the quality of our MSA, we applied the “find-outliers“ https://github.com/veg/hyphy-analyses/tree/master/find-outliers algorithm to our data. Briefly, the script parses a SLAC [Pond and Frost, 2005] results JSON-formatted file, which contains ancestrally reconstructed data of the evolutionary history of the sequences in a phylogenetic tree and iterates over the sequences over a sliding window and examines the number of inferred multiple-nucleotide substitutions and masks problematic codons. This resulted in masking (with “---”) several sites within 34 sequences contained within our alignment, this is done to ensure the quality of our alignment and to limit the spurious effects of molecular sequences from low-quality genome assemblies.

### Phylogenetic inference, annotation, and visualization

For our multiple sequence alignment of Rem2 sequences, we used IQ-TREE v2.2.0.3 [Minh et al, 2020] to perform maximum likelihood (ML) phylogenetic inference. To annotate our phylogenetic tree for various taxonomic groups, we employed the software found at https://github.com/veg/hyphy-analyses/tree/master/LabelTree. To visualize our phylogenetic tree, we utilized the tree viewer available at https://phylotree.hyphy.org [Shank et al, 2018].

### Recombination detection

Genetic Algorithm for Recombination Detection (GARD) [Pond et al, 2006] is a method to screen a multiple sequence analysis for the presence of genetic recombination as a pre-processing step for selection analysis and evolutionary rate inference. The GARD method works by screening an alignment for putative recombination breakpoints and inferring a unique phylogenetic history for each detected recombination block. GARD will search the space of all informative locations for break points in the phylogenetic tree, inferring phylogenies for each putatively nonrecombinant fragment of the tree. Goodness of fit measures are used to determine the number of optimally placed breakpoints by Akaike Information Criterion corrected for small sample size (cAIC) [Hurvich and Tsai (1989)].

### Selection analyses

The multiple sequence alignment and its accompanying unrooted phylogenetic tree was examined with a suite of molecular evolutionary methods, each designed to ask and answer a specific biological question, described below. We performed multiple-test correction via the false discovery rate (FDR) and reported both adjusted and non-adjusted p-values.

All selection analyses were performed in version 2.5.40 of HyPhy [Pond et al, 2020], our set of selection analyses includes the following tests (Table 1):

**Table 1.**
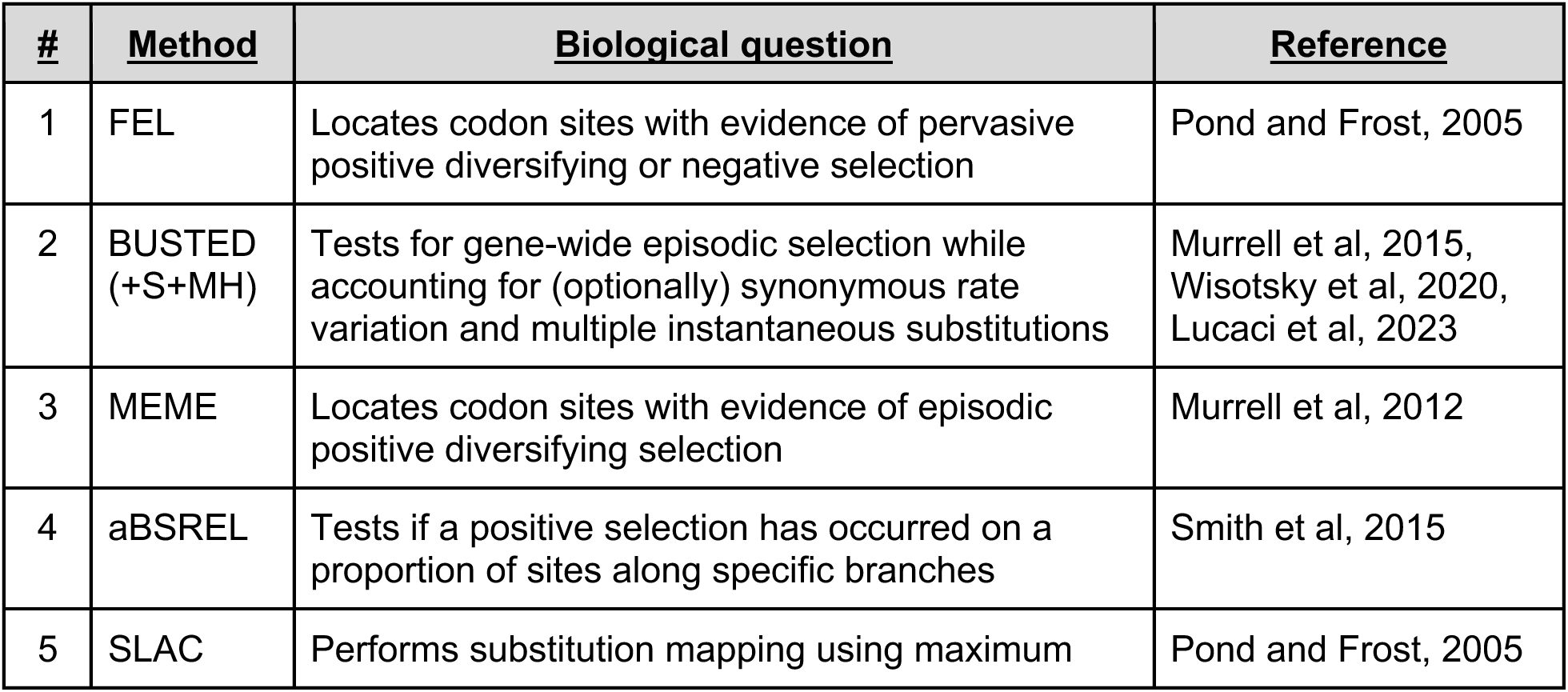
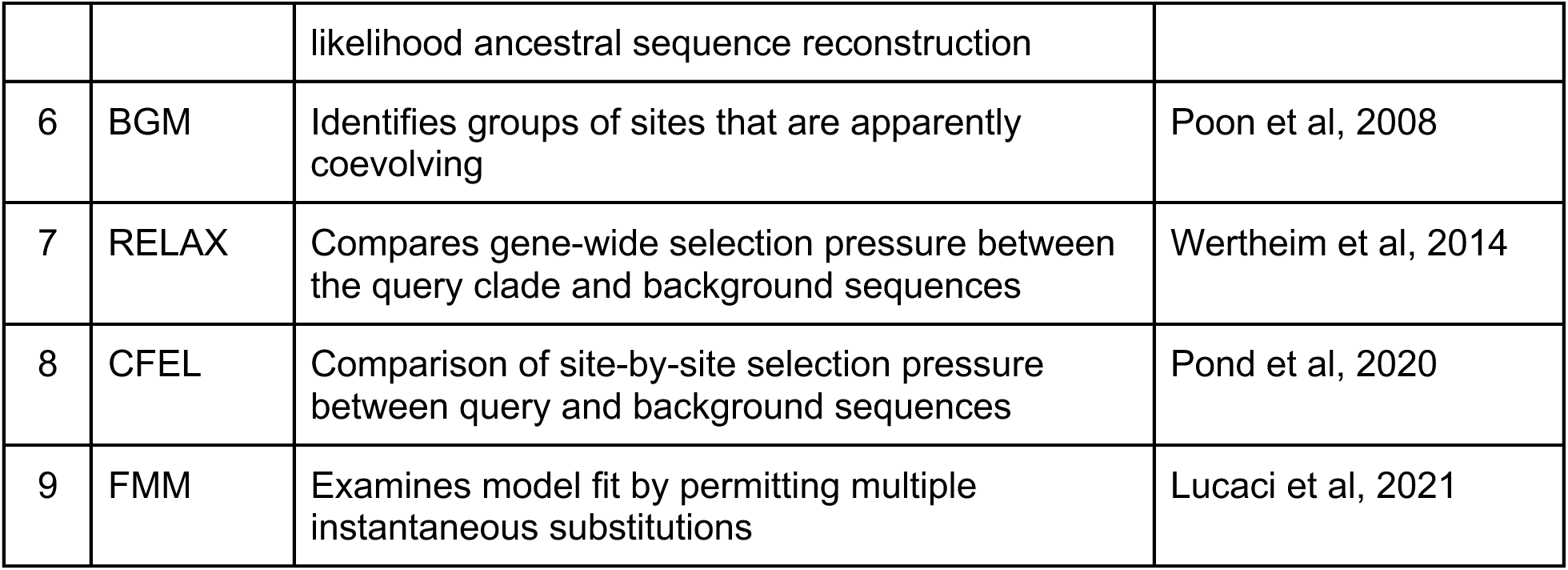
An overview of the selection analyses used in this study. All methods are implemented in the HyPhy software suite.

### <u>A</u>ncestral sequence reconstruction, <u>M</u>olecular <u>E</u>volution, and protein <u>S</u>tructure (AMES)

The AMES analysis is designed to combine ancestral sequence reconstruction of a multiple sequence alignment of extant species with protein structural prediction done with AlphaFold2 [Jumper et al, 2021]. It accomplishes this through the following series of steps. First, we utilize the SLAC JSON output file from HyPhy which is performed for inferring the ancestral state of extinct sequences. The custom Python script “ancestralevolution.py” is used to extract and organize both extant and extinct coding sequences into a multifasta file. Subsequently, the Multifasta file is divided into separate files, each containing a single amino acid sequence per file. Next, we execute the ColabFold tool [Mirdita et al, 2022] available at https://github.com/sokrypton/ColabFold. In batch mode, we utilize the AlphaFold2_batch.ipynb notebook; the tool is executed with default settings, except for enabling the ‘Zip_results’ option. Following the execution, PDB files are extracted, out of the five models used only the best PDB, for each species, with a rank of 1 is selected for further analysis. To compare our results in a combinatorial fashion we ran TM-Align [Zhang and Skolnick, 2005] available at https://anaconda.org/bioconda/tmalign to generate TM-scores, a metric utilized to evaluate the degree of topological similarity among protein structures.

### BUSTED Model testing

We applied the BUSTED model testing and averaging procedure to select the best fitting model for episodic diversifying selection, and to interpret the results of natural selection acting on Rem2. Our goal is to understand which underlying model and its parameters can detect the areas of the dataset which drive the greatest degree of evolutionary signals. Analysis is conducted as a series of experiments in the BUSTED framework of selection analysis in the hierarchical structure and includes the baseline BUSTED method, and extensions to include synonymous rate variation (+S), multiple nucleotide substitutions (+MH), or both (+S+MH). A Snakemake [Mölder et al, 2021] version of the model testing procedure is available at https://github.com/veg/BUSTED_ModelTest.

### Software and Data availability

All software, raw data, and full results, including all HyPhy selection analyzes JSON-formatted result files, used in this study is freely available via a dedicated GitHub repository at: https://github.com/aglucaci/AOC-Rem2. All AMES related software and data used is available at the dedicated Github repository: https://github.com/aglucaci/AMES-Rem2.

## Results

In this study, we estimated the unique selective pressures that have shaped the evolution of mammalian *Rem2* gene across time. Our analysis is based on 175 mammalian species used to construct a multiple sequence alignment (see Table S1 for the full list of species and accessions used). The general structure of the Rem2 protein consists of a proximal disordered region and a distal main functional domain, belonging to the Rem, Rem2, Rad, Gem/Kir (RGK) subfamily of Ras-like GTPases, and is also flanked by another terminal disordered region. We find evidence in 70% of non-invariant codon sites that purifying selection has played a significant role in shaping the evolutionary history of mammalian *Rem2* using the FEL analysis (described in Methods). These results suggest that tight regulation of *Rem2* may be important for maintaining its level of evolutionary fitness. In addition, we also observed 250 out of 626 (or 40%) sites to be invariant at the codon-level, suggesting a highly conserved evolutionary importance for these loci. Evidence of coevolution was also found among 49 pairs of sites, some of which result in complex spatial relationships and potentially lend themselves to epistatic interactions within the protein. Where available, we relate several sites of interest in our results to relevant literature for important considerations for human health.

### Phylogenetic relationship of mammalian Rem2

*Rem2* is an ancient gene with molecular sequences available (at the time of this writing) for *Dipnotetrapodomorpha* which includes tetrapods and lungfish [Hollis et al., 2012]. However, the focus of this study is on Rem2’s preservation throughout the mammalian lineage, where it has been exposed to all significant evolutionary events of the past ∼250-200 million years and has played a role in a diverse and dominant terrestrial animal group [Correll et al, 2008, Edel et al 2010; DeRocher et al., 2014]. This includes all major geological periods including earth impacts (asteroids), oxygenation level changes across time, atmospheric carbon dioxide concentrations, and changes in solar luminosity (radiation flux). An important step to frame the relative relationship of our analysis of mammalian Rem2 is the inference of an appropriate phylogenetic tree (see Figure 1). Our results indicate that even under significant evolutionary pressures over millions of years, that *Rem2* has not been drastically modified in the mammalian lineage or in several of the taxonomic groups contained within *Mammalia* including Glires (rodents and lagomorphs), Eulipotyphla (includes hedgehogs, moles, shrews), Primates, Perissodactyla (odd-toed ungulates), Chiroptera (bats), Artiodactyla (even-toed ungulates), Carnivora. In fact, consistent with other findings [Hutter et al, 2000, Hamel et al, 2006, Anderson et al, 2016, Bennedorf et al, 2022, Lucaci et al, 2022], *Rem2* has alignment characteristics consistent with ancient genes that tend to be highly conserved and therefore are more likely to exhibit an abundance of purifying selective changes as opposed to severe adaptive evolutionary changes [Karczewski et al, 2020]. Indeed, genes that undergo strong purifying selection, exhibiting minimal evolutionary changes, are more prone to being associated with severe human diseases [Lek et al, 2016]. While we examined selection pressures between taxonomic groups using the RELAX and Contrast-FEL methods, we did not find statistically significant results. This could be explained by the similar biological roles and fitness effects that Rem2 has in our groups under study. Using the aBSREL method, we found evidence of episodic diversifying selection in 3 out of 332 branches (results not shown) in our phylogeny of mammalian *Rem2,* specifically for the following organisms: *Equus przewalskii*, *Colobus angolensis*, and *Capra hircus*.

**Figure 1.**
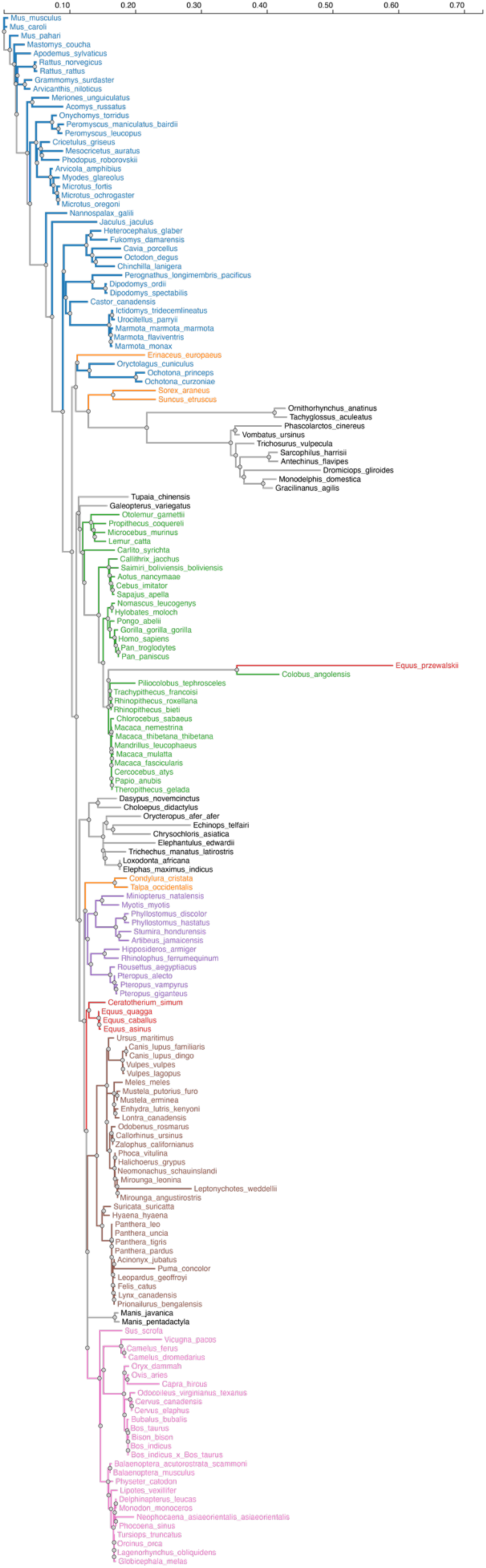
Phylogenetic relationship of the Rem2 gene in *Mammalia*. The evolutionary history of Mammalian Rem2 was inferred using the maximum likelihood (ML) method via IQ-TREE (see additional details in the Methods section). The inferred tree is shown and is drawn to scale (expected substitutions/site). We annotated our inferred phylogenetic tree with major taxonomic groups (if more than three species were contained in a single taxonomic group) from Mammalia including: Glires (Blue, rodents, and lagomorphs), Eulipotyphla (Orange, includes hedgehogs, moles, shrews), Primates (Green), Perissodactyla (Red, odd-toed ungulates), Chiroptera (Purple, bats), Artiodactyla (Pink, even-toed ungulates), Carnivora (Maroon). Some branches (in black) are unlabeled and were not used in across-taxa comparative analyses, discussed below.

### Evidence of gene-wide evolution in Rem2

The conservation observed in the nucleotide sequences among mammalian *Rem2* and its various taxonomic subgroups, is indicative of a strong selective pressure acting on the *Rem2* gene. To examine evidence for gene-wide episodic diversifying selection (EDS), we used the BUSTED+S (branch-site unrestricted statistical test for episodic diversification, with synonymous rate variation) method on our multiple sequence alignment and inferred phylogenetic tree, which statistically tested three omega rate categories while integrating over their assignments to all branches and sites. We observe a statistically significant result (averaged likelihood ratio test (LRT) p-value 0.0075 ≤ 0.05) which provides evidence of whole gene and branch-site level EDS in the Mammalian Rem2 gene. The ω_3_ rate (which is the measure of adaptive evolution in this method) was >100. and occupied a proportion of 0.06% of the data (Table 2). This evidence suggests that a minute fraction of (branch-site) pairs in this gene are driving the signal for adaptive evolution. While this branch-site method revealed the occurrence of EDS within the Mammalian *Rem2* gene, and is supported by suggestive evidence, based on the evidence ratio (ER), a heuristic for evolutionary signal in this method based on site-level likelihood ratios [described in Murrell et al, 2015], in 15 sites distributed broadly (Figure S1), with an Evidence Ratio (ER) threshold of 10, across the *Rem2* locus, it does not provide conclusive evidence of site-level selection properties. For this we will carry out a more sensitive site-level analysis below.

**Table 2.**
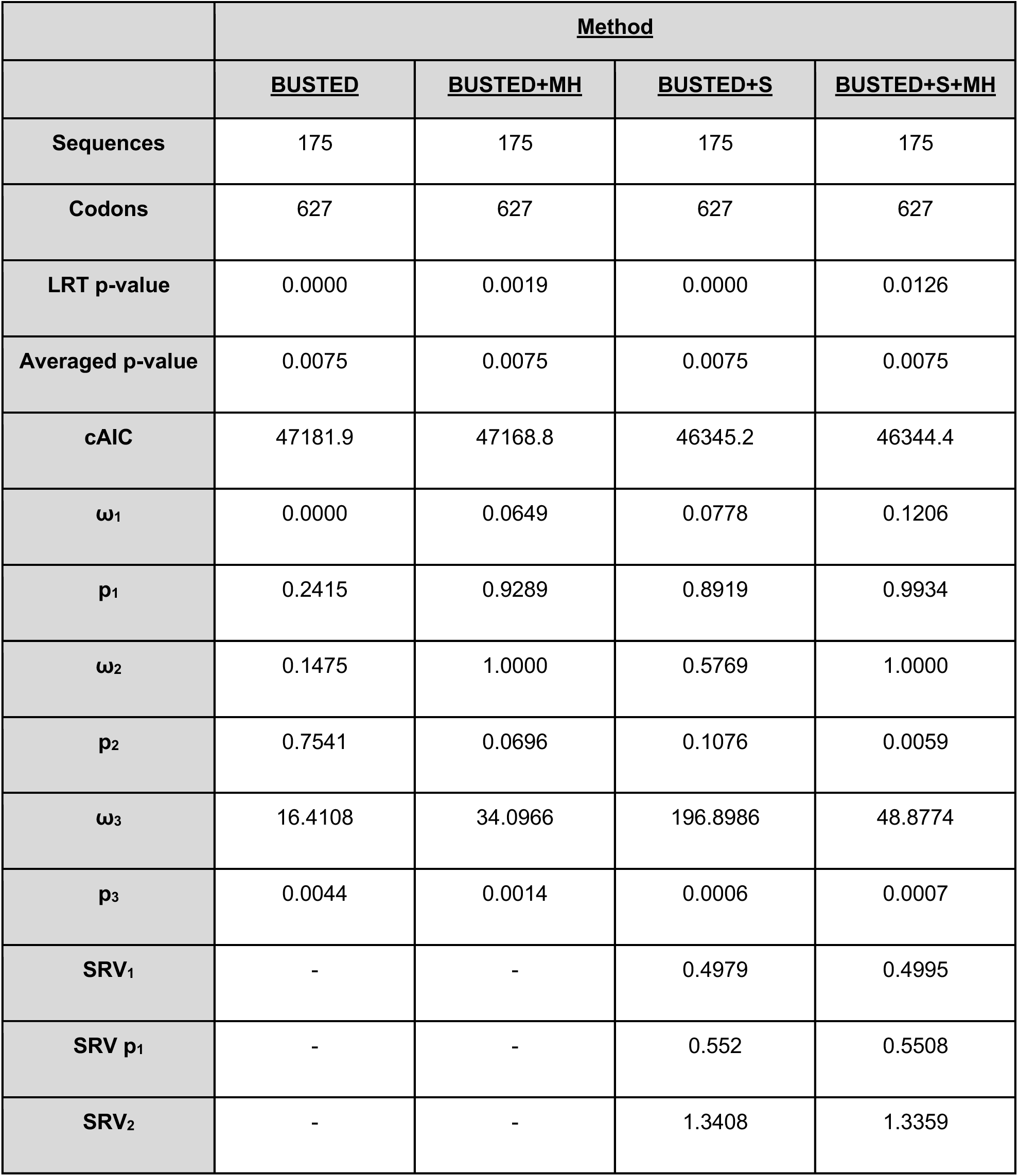
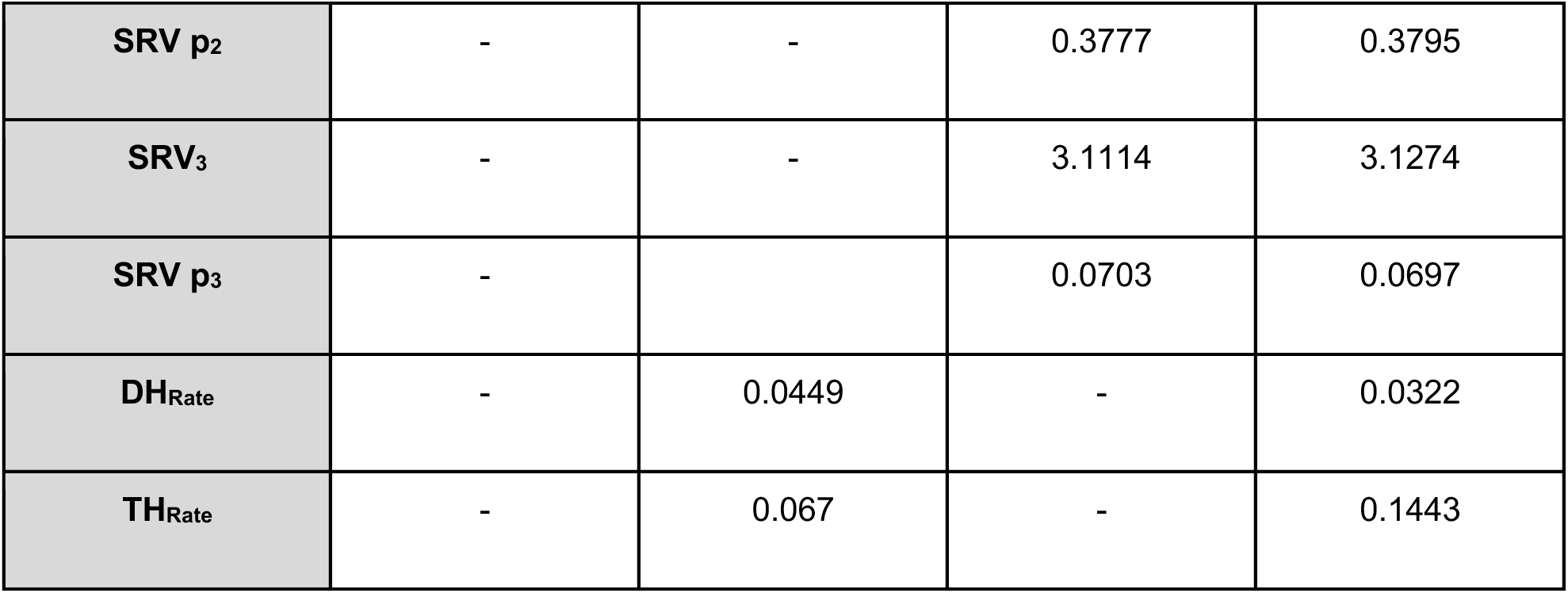
Evidence of Gene-wide evolution in Mammalian Rem2. This table describes summary statistics for the BUSTED ModelTesting approach. Briefly, in each column we applied one of four methods, each of which either accounts for a biological phenomenon of synonymous rate variation (+S) and/or multi nucleotide (+MH) substitutions or does not. Sequences and Codons correspond to the number of taxa and codon sites in the alignment, respectively. LRT p-value and Averaged p-value correspond to the unadjusted and adjusted (see Methods for details). cAIC is the small-sample Akaike information criterion. Omega rates 1-3 correspond to inferred omega estimates, while “p” 1-3 values are the proportion of the data that these rates were found to be the best fit. SRV rates 1-3 correspond to inferred synonymous rate variation estimates, while “SRV p” 1-3 values are the proportion of the data that these rates were found to be the best fit. DH and TH rates are the inferred double nucleotide and triple nucleotide substitution rate estimates.

|  | <u>Method</u> |  |  |  |
| --- | --- | --- | --- | --- |
|  | <u>BUSTED</u> | <u>BUSTED+MH</u> | <u>BUSTED+S</u> | <u>BUSTED+S+MH</u> |
| **Sequences** | 175 | 175 | 175 | 175 |
| **Codons** | 627 | 627 | 627 | 627 |
| **LRT p-value** | 0.0000 | 0.0019 | 0.0000 | 0.0126 |
| **Averaged p-value** | 0.0075 | 0.0075 | 0.0075 | 0.0075 |
| **cAIC** | 47181.9 | 47168.8 | 46345.2 | 46344.4 |
| **$\omega_1$** | 0.0000 | 0.0649 | 0.0778 | 0.1206 |
| **$p_1$** | 0.2415 | 0.9289 | 0.8919 | 0.9934 |
| **$\omega_2$** | 0.1475 | 1.0000 | 0.5769 | 1.0000 |
| **$p_2$** | 0.7541 | 0.0696 | 0.1076 | 0.0059 |
| **$\omega_3$** | 16.4108 | 34.0966 | 196.8986 | 48.8774 |
| **$p_3$** | 0.0044 | 0.0014 | 0.0006 | 0.0007 |
| **SRV<sub>1</sub>** | - | - | 0.4979 | 0.4995 |
| **SRV $p_1$** | - | - | 0.552 | 0.5508 |
| **SRV<sub>2</sub>** | - | - | 1.3408 | 1.3359 |

**Table 2. Evidence of Gene-wide evolution in Mammalian Rem2.**
|  |  |  |  |  |
| --- | --- | --- | --- | --- |
| **SRV p<sub>2</sub>** | - | - | 0.3777 | 0.3795 |
| **SRV<sub>3</sub>** | - | - | 3.1114 | 3.1274 |
| **SRV p<sub>3</sub>** | - | - | 0.0703 | 0.0697 |
| **DH<sub>Rate</sub>** | - | 0.0449 | - | 0.0322 |
| **TH<sub>Rate</sub>** | - | 0.067 | - | 0.1443 |

Multi-nucleotide substitutions involve changes in multiple adjacent nucleotides in DNA and may have significant impacts on gene evolution by creating novel genetic variation, increasing the accessibility of evolutionary pathways, and altering the structure and function of proteins. Studies have shown that multi-nucleotide mutations can contribute to adaptation and evolutionary innovation in a range of organisms [Steward et al 2022, Hensley et al, 2021, Cohen et al 2020, Silva et al, 2023]. Considering these studies, we have applied a recently developed selection analysis [Lucaci et al, 2023] that incorporates two confounding biological processes: synonymous rate variation (SRV) and multi-nucleotide substitutions (MH) into the BUSTED framework. The overall result still finds evidence of selection in the mammalian Rem2 gene, raising our confidence that the MH-inclusive and traditional BUSTEDS analysis agree. However, we observe a lower ω_3_ rate value of ∼50 over a slightly smaller proportion (0.07%) (Table 2) of the data, indicating that while some of the evolutionary signal across sites is explained by the MH parameters (we estimate non-zero values for the MH rates), we retain statistical significance. Our interpretation is that, overall, based on our model averaging approach (described in Lucaci et al, 2023), our results demonstrate evidence suggesting that mammalian Rem2 is under episodic diversifying selection.

### Pervasive purifying selection in mammalian Rem2

Our analysis of negative selection in mammalian *Rem2* was carried out using the FEL method (see Methods section for details) in HyPhy. The results (see Figure 2) provide a measure of the extent of purifying selection for each site in the human *Rem2* gene and indicate that 70% of non-invariant codon sites show a statistically significant estimate of purifying selection. To visualize the spatial organization of our results, the dN/dS estimates for the entire alignment were plotted with a confidence interval of the 95% lower and upper-bound estimates (see Figure 2 or Table S2). Overall, mammalian *Rem2* exhibits broad evidence of purifying selection in proximal disordered regions and in distal RGK and disordered regions. These results suggest that purifying selection has played a predominant role in shaping the functional and mature *Rem2* gene, which exhibits remarkable conservation across evolutionary epochs, leading to its rapid evolutionary adaptation in other genes and taxa.

**Figure 2.**
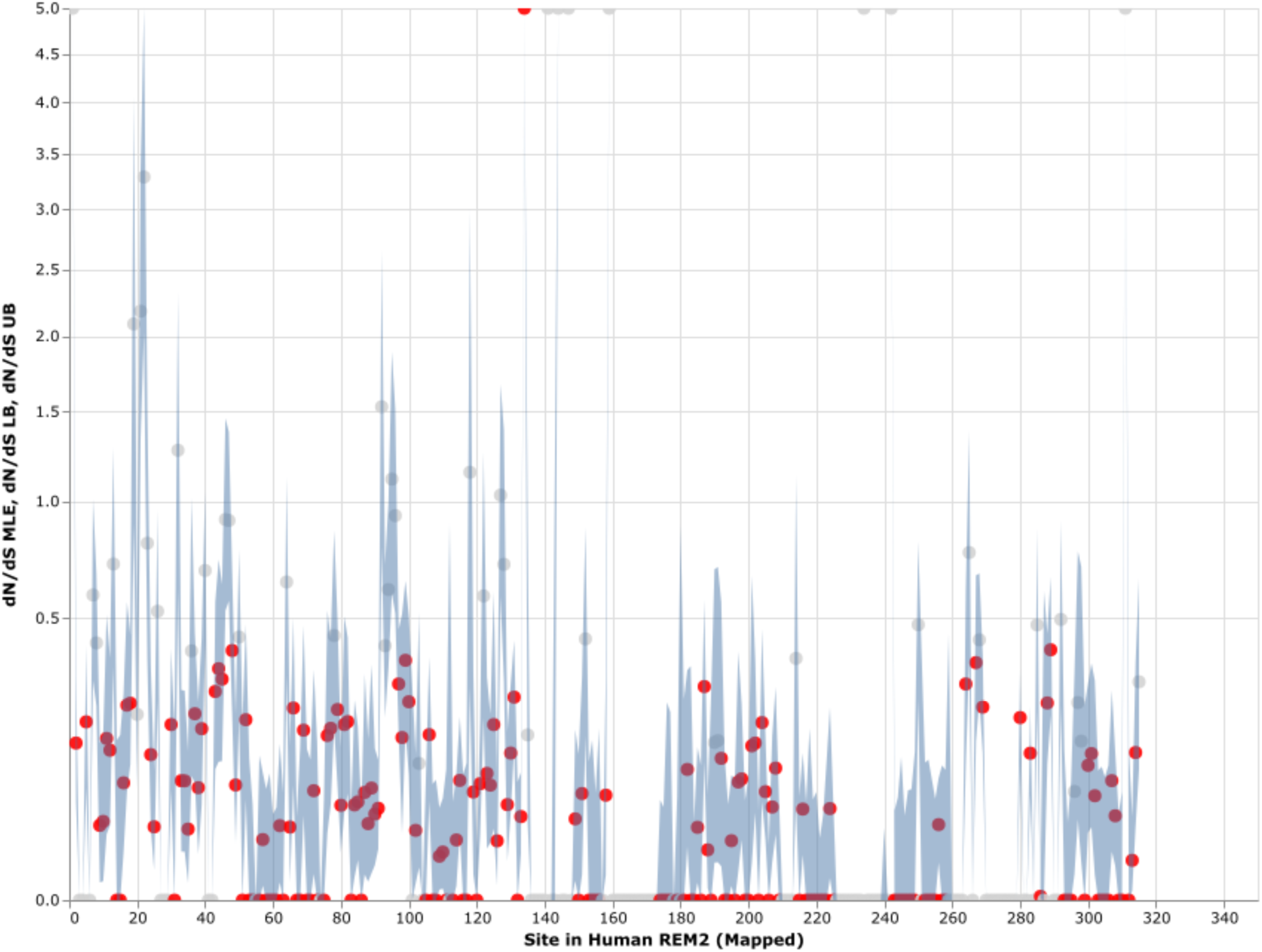
Strict purifying selection across sites in the Mammalian *Rem2* gene. The FEL analysis, used to detect sites of pervasive positive or negative selection, reveals statistically significant sites in the Mammalia Rem2 gene alignment. Sites that are statistically significant (adjusted LRT p-value ≤ 0.1, for FDR) are indicated in red. In this chart, we plot the estimated values of omega (or dN/dS, Maximum Likelihood Estimate (MLE)) for each site in the alignment (non-statistically significant sites are plotted in gray). Additionally, we plot 95% confidence intervals (CI) for the omega estimate for each site (light blue).

While we searched for evidence of adaptively evolving sites using the FEL (Figure 2 and Table S2), we found a single positively selected site using FEL, after multiple tests correction, corresponding to alignment codon site 170 (human *Rem2* site 134). In comparison, using the MEME (Table S3) method, after applying multiple tests correction, we found no sites remained statistically significant. We have included the MEME table of results for completeness in the supplementary material. As more information becomes available (i.e., number of mammalian genomes, or increasing the scope of the study to examine taxonomic groups outside of mammalian) we expect that some of these sites may become statistically significant, indicating a role for adaptation to unique environmental pressures.

Purifying selection plays a crucial role in shaping the evolution of genes in mammals by maintaining functional genes, removing harmful mutations, and preserving genetic diversity. By analyzing orthologous sequences across species, we observe the broad distribution of negatively selected sites across the *Rem2* gene (see Figure 2, Table S2). Protein-coding sequences with highly constrained structures are expected to fix nonsynonymous mutations at a slower rate due to the maladaptive nature of changes such as what we observe with negatively selected sites across *Rem2*.

### Broad evidence of coevolutionary forces within Rem2

To examine the coevolution of codon sites in mammalian Rem2, (i.e., if a particular codon site was evolving in a relationship with another codon) we used the Bayesian graphical models (BGM) method. This method infers the substitution history of an alignment with maximum-likelihood ancestral sequence reconstruction and maps these to the phylogenetic tree, which allows for the detection of correlated patterns of substitution. In our results we find evidence for pairs and groups of 49 putatively coevolving sites in our mammalian *Rem2* alignment (see Figure 4, and Table S4). This data suggests that interaction dynamics may be impacted in tertiary space of the folded protein These results may also be evidence that coevolving sites may be related to other fitness consequences (i.e., offering compensatory mutations) for maladaptive changes in another part of the protein sequence.

**Figure 4.**
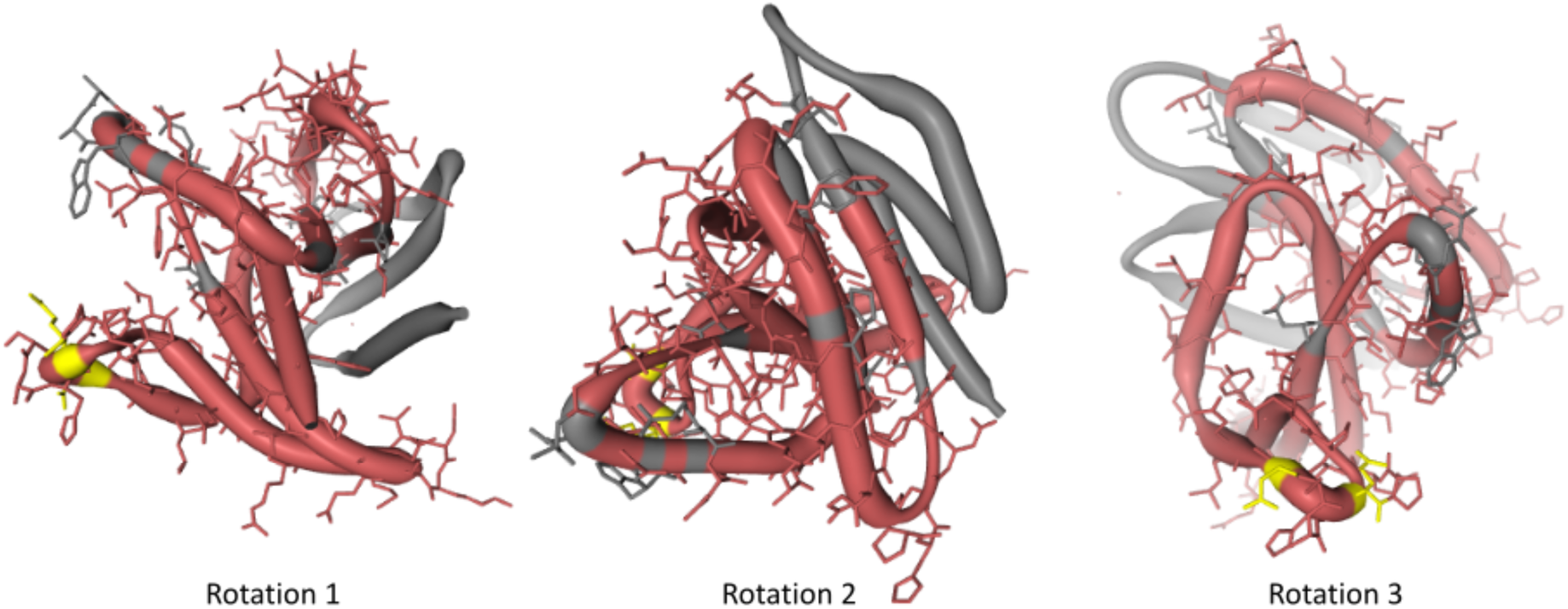
Protein structural analysis of Human Rem2 to highlight selected sites in the conformational space. In this figure, we demonstrate the structural configuration of the Human Rem2 protein (https://www.rcsb.org/structure/3CBQ) with selected sites from our results. This conformation represents the crystal structure of the Human Rem2 GTPase with bound GDP where we highlight purifying (pink) evolving sites, with neutral sites represented in gray and invariable sites highlighted in yellow. We include several rotations with arbitrary degrees to highlight the view of these sites in the folded protein space. This figure was generated in an interactive ObservableHQ [Perkel, 2021] notebook that is rotatable in the 3D space, and is available here: https://observablehq.com/@aglucaci/structure-rem2. The PDB structure is limited to the functional domain of Rem2 which limits our ability to highlight all our sites of interest (SOI), therefore we have limited our annotation only to the modeled sites in the structure.

When we reviewed these sites, we noticed that several pairs of coevolving sites (Figure 4) occurring only in the human Rem2/RGK (Table S4), and some pairs with one site in the proximal disordered region and another within the human RGK region, linking these distant regions. Several regions display complex geometric features including a network of interactions spanning multiple sites: 86, 604, 72, 68, 120. When mapped to the human *Rem2* reference sequence all these sites fall within the RGK region, and correspond to 64, 319 (in RGK), 52, 48, 98, respectively. When we examined the gnomAD database [Karczewski et al, 2020] for the clinical relevance of these sites, we found the following human variants at these sites:

Disruption of the delicate balance maintained in protein evolution of this type, in this network or any of the coevolving networks observed in mammalian *Rem2* could have fitness altering clinical relevance and are worth further investigation. We appreciate that sites 48, 64, 98 in human Rem2 are in a coevolutionary network together and have possible functional implications when mutated (Table 3).

**Table 3.** Clinical implications of coevolutionary important codon sites in Rem2. In this table we denote the consequences of mutations found in the gnomAD database for codon sites where we found evidence of coevolutionary forces. Note that mutations at positions 23,353,970 and 23,353,971 in Rem2 fall within a potential MNV (*) where the variant is found in phase in 1 individual, altering the amino acid sequence, the combined change is as follows: aGA to aTT becomes a R to I change.

| <u>Codon Site in Human Rem2</u> | <u>Genome Position (hg19)</u> | <u>Variant Type</u> | <u>Nucleotide change</u> | <u>Amino acid change</u> | <u>Mutation Consequence</u> | <u>PolyPhen</u> | <u>SIFT</u> |
| --- | --- | --- | --- | --- | --- | --- | --- |
| 48 | 23,353,918 | Deletion | CTG->Cdel | Leu48GlyfsTer37 | Frame shift | - | - |
| 64 | 23,353,969 | SNV | A->T | Arg64Ter | Stop gained | - | - |
| 64 | 23,353,970 | MNV* | G->T | Arg64Ile | Missense | Possibly damaging | Deleterious |
| 64 | 23,353,971 | MNV* | A->T | Arg64Ser | Missense | Possibly damaging | Deleterious |
| 98 | 22,884,863 | SNV | C->T | Ser98Phe | Missense | Probably damaging | Deleterious |

### Structural view of selected sites on the Human Rem2 protein

To gain an appreciation of the spatial distribution of our sites of interest returned by our statistical methods we relied on a PDB structure to generate a three-dimensional reconstruction. We examined the crystal structure of the human Rem2 GTPase with bound GDP with a modeled residue count of 169. We focus on the main functional domain of the *Rem2* gene and its function as a GTPase as an example of spatial protein-ligand interactions. We believe that this additional layer of biological and physical-chemical interaction has contributed to the complex evolution of the functionality of Rem2. By mapping our results from the selection analyses listed above, we examined sites with evidence to be under the effect of a purifying selective regime in the evolutionary history of Rem2 in mammals.

Although the immediate effects of evolution on protein structure (Figure 4) are not well understood and cannot be fully verified computationally but rely on experimentally determined results, they have implications for our understanding of Rem2 neurobiology. Our coevolutionary and selection analyses suggest that the RGK, a region in Rem2 known for its functional effects, is implicated. Our 3D modeling (based on a known PDB structure) indicates that the patterns and underlying networks of coevolving sites are in complex relationship with each other, which may have undiscovered functional impacts. Therefore, we predict that evolutionary changes in Rem2 are likely to reflect some form of specialization, divergence in function, or interaction with protein partners at different stages in its evolution in mammals. Further research may reveal additional connections that can provide deeper insight into Rem2’s diverse functions and its role in human health and disease.

### Ancestral sequences of mammalian Rem2 reveal evolutionary dynamics and conservation

We used ancestral sequence reconstruction (ASR) and predicted protein structure to shed light on the evolutionary history and conservation of mammalian Rem2. By analyzing 175 extant and 156 reconstructed extinct sequences, our results suggest some degree of structural difference across Rem2 in the core functional RGK domain and in its intrinsically disordered regions, highlighting the complex interplay between purifying selection at the sequence level (noted above) and protein structural evolutionary trajectory (Figure 5). Sequence-level conservation of Rem2 across different mammalian species helps support its functional importance and potential roles in various biological processes (Figure 6). Our approach also belies the importance of examining the evolutionary history of ancient genes and pathways [Goulty et al, 2023].

**Figure 5.**
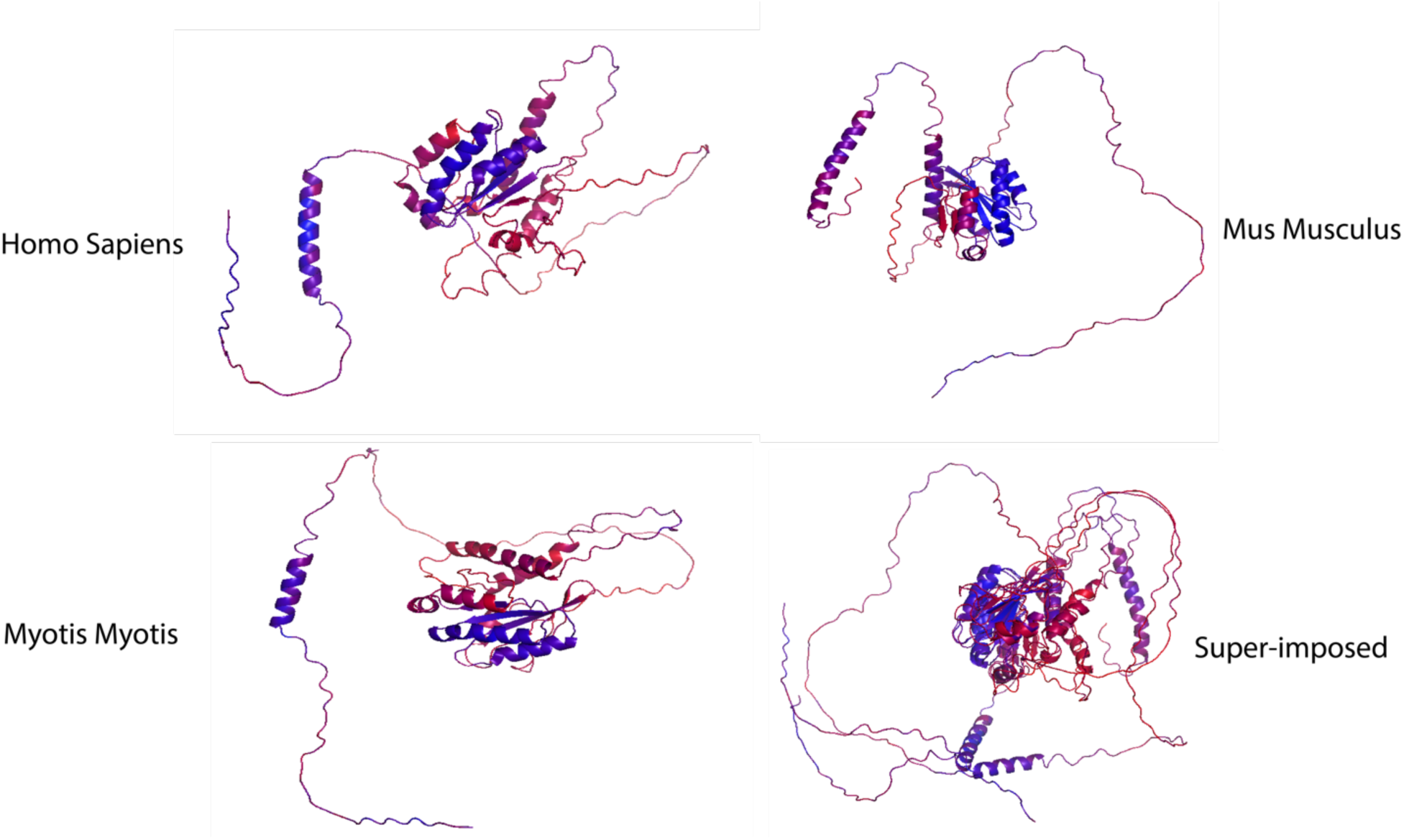
Ancestral reconstruction of predicted protein structures reveals an evolutionary conserved core functional domain in RGK. We constructed the ancestral sequences of Rem2 to unravel its evolutionary trajectory. The color spectrum is as follows: red (low confidence sites) to blue (high confidence sites). Additional details about this method (AMES) are provided in the Methods section. Root Mean Square Deviation (RMSD), TM-score is a quantitative measure to evaluate the degree of topological similarity between protein structures. Protein structures were rendered in PyMOL version 2.5.5.

**Figure 6.**
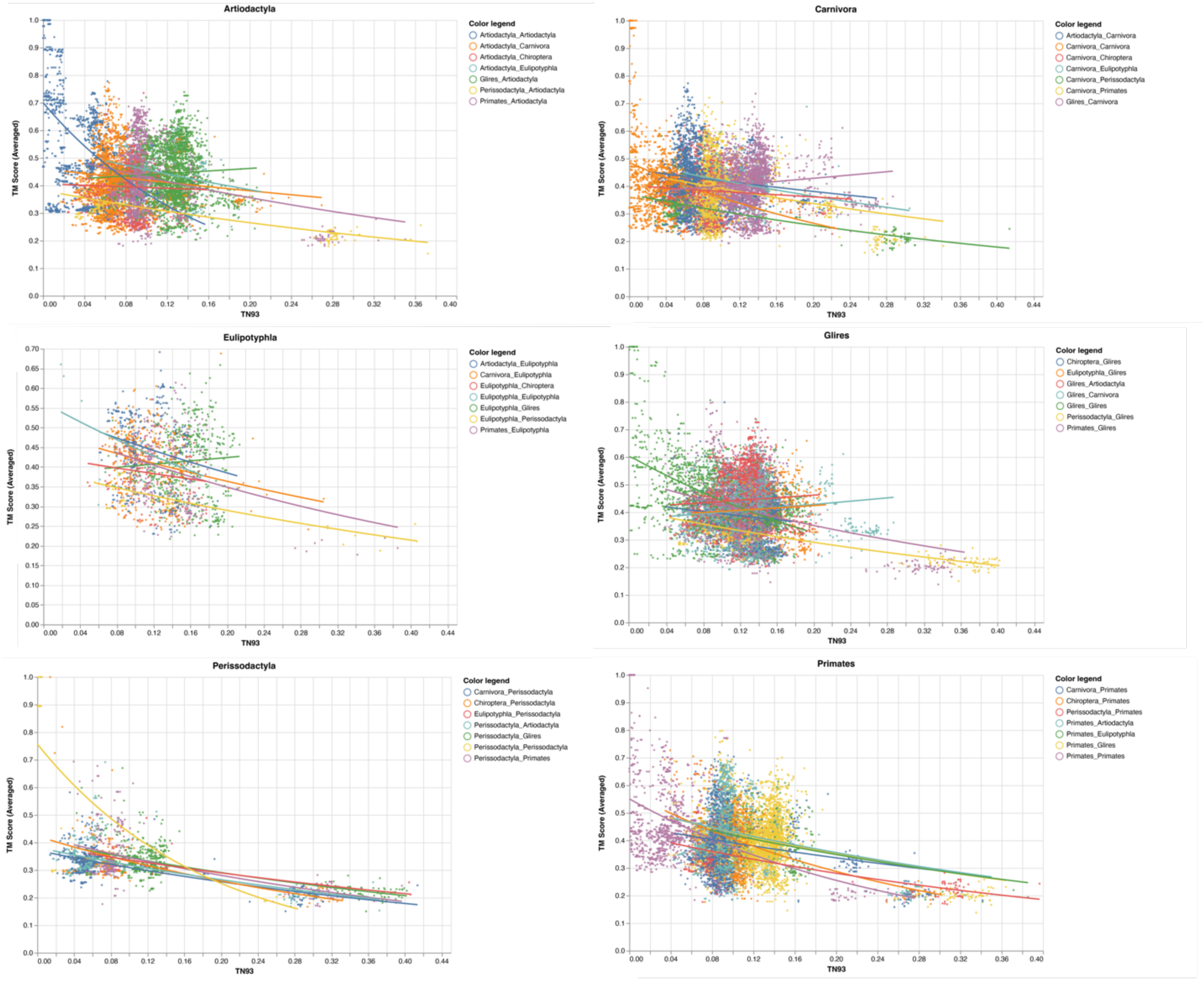
A comparison of genetic and structural differences in mammalian *Rem2.* In this chart we show scatterplots and regression lines for our measures of genetic distance (via TN93) and structural distances (via TM-Score calculations), each subplot is faceted by the clade of interest, the color legend defines the taxonomic annotation of each pairwise comparison.

However, how it accomplishes this in different species may be slightly different at the protein-structural level. RMSD (Root Mean Square Deviation) is a measure that quantifies the average distance between atoms, typically the backbone atoms, of proteins that have been superimposed. When examining the results for our RMSD calculations we find an average score of 4.95 (std 0.82, Table S5), indicating a level of discordance across the range of structures; lower values of RMSD are better, with a score of <2 considered in good agreement.

To circumvent some of the short-comings of RMSD we also utilized the TM-score, a metric that prioritizes smaller distance errors over larger ones, making it more sensitive to global fold similarities rather than local structural variations and has a length-dependent scale to normalize the distance errors making the magnitude of the TM-score length-independent for random structural pairs. When examining the results for our TM-score (using TM-align) calculations we find an average score of 0.38 (std 0.11, Table S5), also indicating a level of discordance across the range of structures with values above 0.5 considered in about the same structural fold and values below 0.3 considered to be random (Figure 6). When filtering for strong pairwise TM-score’s (above 0.5) we find that 10.5% of our dataset demonstrates protein structural similarity (Table S5). However, our results are not to be overinterpreted and may be largely confounded by low confidence structural inference (low pLDDT scores), difficulty in aligning intrinsically disordered regions (IDR) of Rem2, or alignment errors. It may also be useful to examine Rem2’s protein structural domain evolution, including instances where specific domains are gained or lost across the phylogenetic tree and how these impact the comparison of structural inference.

Comparison of molecular (TN93 genetic distance) and structural distances (TM-Score) in Figure 6 reveal a general trend of agreement between increased genetic distance and increased structural changes. However, we note that for even small genetic distances (<0.05) we observe a large variance in both RMSD and TM-Score, indicating that divergence could be driven by either the disordered region of the Rem2 protein or by neutral mutations in its core functional domain, and convergence explained through conservation of sequence and protein structure for closely related species. Taken together molecular sequence divergence across *Mammalia* could spur evolutionary innovation in *Rem2*.

## Discussion

In the present study, we collected all sequence data available for the orthologous *Rem2* gene, gathering from a diverse set of species within the *Mammalia* taxonomic group. Our results indicate that unique evolutionary processes have shaped the evolutionary history of *Rem2*. Our study consisted of performing several evolutionary analyses that each asked and answered specific biological questions (see Methods section for additional information) to quantify the signals of natural selection in our dataset. Our results revealed that *Rem2* is under tight purifying selection (Figure 2) distributed broadly across the gene, suggesting its sequence and structural conservation is important in maintaining specific protein functionality. We also identified novel substitutions and areas of interest (see Figure 2, Table 3, Figure 3, Table S2, and Table S3) across regions of *Rem2* that may provide potential targets for designing molecular therapeutic approaches with significant impact on protein function. We also observe a complex network of spatial coevolution within this gene, with evidence that 49 pairs of sites have evolved co-dependently (Figure 3, Table S4).

**Figure 3.**
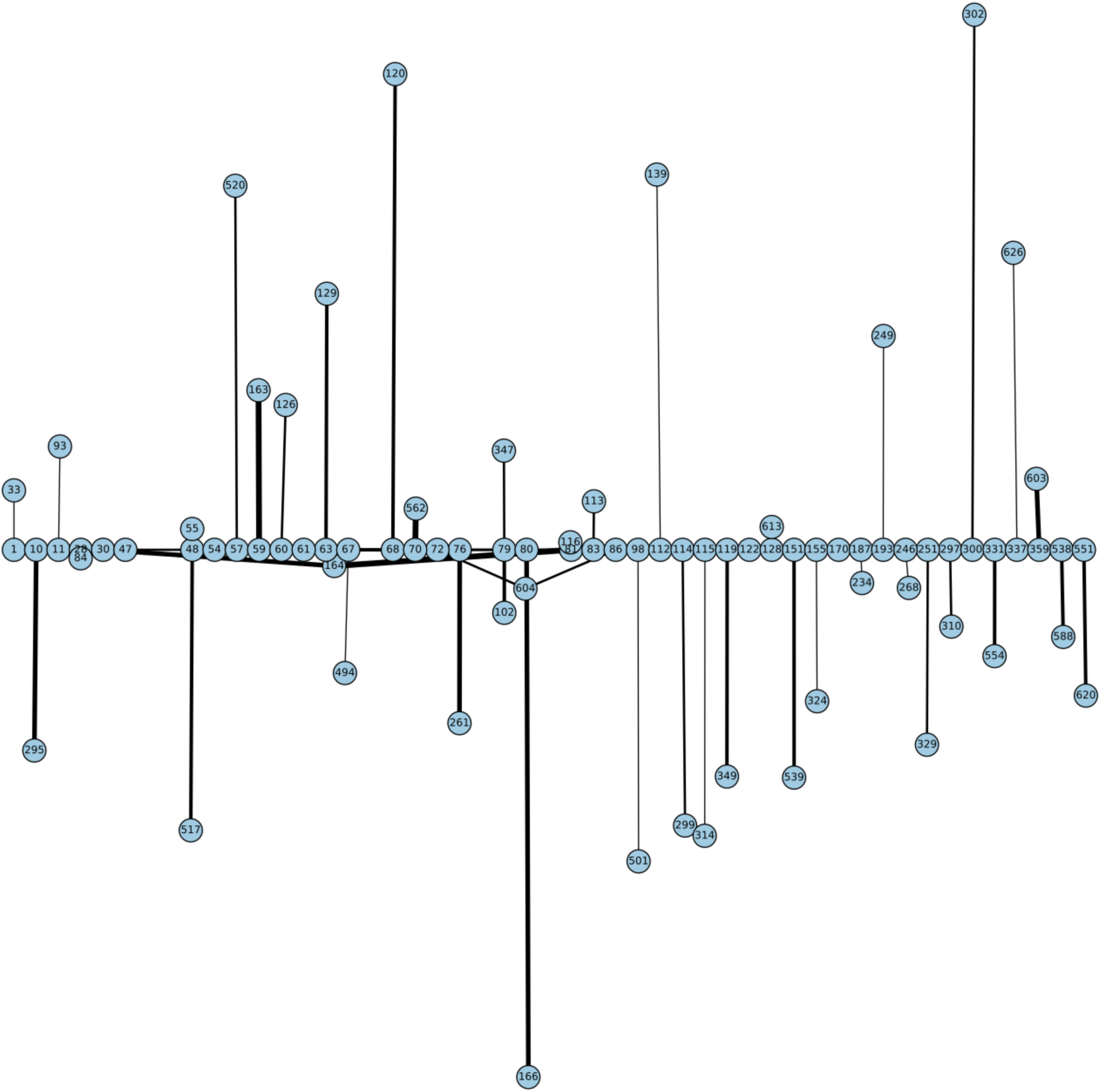
The Bayesian Graphical Modeling (BGM) analysis, a method to detect co-evolving sites in a gene, of Mammalian Rem2 found 49 pairs of coevolving sites out of 627 total sites, to be statistically significant with a posterior probability threshold of 0.5 (see Table S4). In this chart, we plot the statistically significant coevolving pairs (nodes) with the number of shared substitutions between pairs of coevolving sites (edges) controlling the thickness and color of the line, with larger connections indicating a higher number of shared substitutions. The extent of coevolution tends to be distributed both proximally and distally across the *Rem2* gene, indicating interaction between the early region and its conserved core functional domain. Which suggests that sequence variation incurred in this region most likely has a deleterious impact on protein fitness. Note that these sites have been mapped to correspond to sites in the Human Rem2 protein.

### Abundant purifying selection across the Rem2 gene in Mammalia

Our results demonstrate a high degree of purifying selection observed across *Rem2*, which we hypothesize is based on the critical role of Rem2 on its underlying network of genes governing homeostasis and normal brain development [Burkhardt et al, 2023]. We believe that the conservation of function, and gene family evolution plays a role in shaping the evolutionary history of Rem2 based on the maintenance of functional integrity. This interpretation is also consistent with the observation that *Rem2* plays an important role in nervous system development [Ghiretti et al., 2011, 2014; Moore et al., 2013, 2018; Edel et al., 2010].

### No strong evidence of diversifying evolution at specific sites in the Rem2 gene in Mammalia

*Rem2* has been recognized as a crucial gene for synaptogenesis and the regulation of dendritic morphology [Ghiretti and Paradis 2011; Ghiretti et al., 2014]. However, due to significant evolutionary pressure shaping *Rem2* in novel environments we expected to find a small number of sites evolving adaptively. Unfortunately, new sites under positive diversifying selection were not found in the *Rem2* gene (see Table S3) after multiple tests correction. By observing the diversification of amino acid changes which can drive species-specific adaptation and/or increase function in relation to environmental pressures, helping the scientific community understand the evolution of *Rem2* in a closer species-specific manner. However, with a limited number of species in our Rem2 data, our results suggest that future research may yield results that hold important functional properties and may play a role in organismal disease. We expect that increasing the taxonomic range under study to all available sequences may yield different results. Additional research is needed to determine the significance of these adaptively evolving sites, including specialized or regulatory functions and their relation within the network of genes necessary for neural cell diversity and development. The 23 sites we found within our MEME results with statistically significant unadjusted p-values (Table S3) may be a good starting point. While the clinical and functional significance of these sites remains a hypothesis, this study highlights their potential importance, and we look forward to further testing in controlled laboratory conditions.

### Discovery of coevolving pairs in the Rem2 gene in Mammalia

We found a significant number of putatively coevolving sites within the *Rem2* gene (as shown in Figure 3 and Table S4), which represents a new aspect of Rem2 biology not reported before. Here, the coevolution of codon sites refers to a phenomenon where substitutions at one codon site co-occur along the same branches as changes occurring at another codon site. Multiple factors contribute to the coevolution of codon sites within a protein, such as functional or structural constraints, interactions at the amino acid or mRNA level, and selective pressures. For example, a scenario where specific residues hold importance for compensatory changes which may play a crucial role in maintaining a protein’s proper structure and function especially in core functional domains. Coevolving sites were not limited to a particular domain or motif and were distributed throughout the entire Rem2 gene broadly, with some interactions in the RGK functional domain region. However, there were also coevolving sites in regulatory regions (outside of the RGK domain) linked to RGK domain sites. These relationships may play a significant role in shaping the evolution of this crucial gene and its enzymatic properties. The new evidence of coevolution suggests the importance of these sites in regulating the functional domain of Rem2. These residues may form crucial interactions for the functional integrity of Rem2, and the specific pairs that span the Rem2 RGK and non-RGK regions suggest a new mechanism by which the functional domain may regulate the RGK or vice versa. Alternatively, these coevolving pairs are part of a network of residues adapting to new environments post-speciation events and therefore serve species-specific functional requirements.

### Potential protein structural implications of evolving sites

Timothy syndrome is a rare genetic condition that is caused by mutations in the gene encoding the calcium channel protein CACNA1C. Recent studies have shown that the Rem2 gene is involved in modulating the function of CACNA1C, and thus may play a role in the pathogenesis of Timothy syndrome, as has been observed with Gem (see above references). In Timothy syndrome, mutations in the CACNA1C gene result in a gain-of-function in the calcium channel, leading to an increase in intracellular calcium levels. The increased calcium levels can cause a range of symptoms, including heart problems, developmental delays, and autism. Studies have shown that the Rem2 protein can interact with CACNA1C and modulate its activity, suggesting that the *Rem2* gene may be involved in the pathogenesis of Timothy syndrome. However, more research is needed to fully understand the relationship between *Rem2* and Timothy syndrome.

Our AMES analysis can also serve as a resource for the scientific community interested in Rem2’s evolutionary trajectory. The dataset provides utility for examining additional hypotheses involving the properties of molecular sequences and the protein structural dynamics. It may be particularly interesting to examine where in evolutionary time specific functional domains were lost or gained in Rem2 in response to species-specific neurodevelopmental needs. We envision that approaches could be applied for structural-functional prediction and can be coupled with datasets from carefully designed experimental projects in future studies.

### Limitations of our computational evolutionary analysis

This study focused on Rem2 sequences from the Mammalia taxonomic group instead of examining a more inclusive dataset for Rem2 sequences from all of Dipnotetrapodomorpha (lobe-finned fishes including tetrapods and lungfishes). Therefore, while our selection analyses are applicable to mammals, our results do not capture the entirety of Rem2’s evolutionary history (Puhl et al, 2014). We also do not explore the mutational patterns occurring outside of protein CDS (i.e., UTR’s) which include complex structure and dynamics at the mRNA level of non-coding regions in the *Rem2* gene. An additional limitation of the current analysis is due to the presence of indels, spatially distributed across several regions of the Rem2 alignment. Although there is a risk that particularly gappy regions of our multiple sequence alignment may be a computational artifact of the alignment procedure or from the quality of the genomic assembly procedures, based on all other outputs we believe that our results are reasonably interpreted and have subsequently tolerated these potential effects.

## Conclusions

Our research modeled the natural history of changes in the Rem2 gene across 175 mammalian genomes spanning about ∼200 million years of evolutionary history. By analyzing a deeper phylogenetic dataset, one could reveal additional information on the substitutional history and lineage specific adaptations in structure and function of this gene. Overall, we believe that experimental techniques in biology can benefit from these types of evolutionary analysis, specifically where bioinformatics analysis can provide valuable information to guide experimental investigation and increase the efficiency and accuracy of biological research [Cisek et al, 2021]. Importantly, evolutionary history can inform both predictive modeling about the outcomes of mutations in individuals and the integration of multiple levels of information can lead to novel insights for Rem2. In addition, both experimental design for fitness-altering regions of Rem2 and target identification for chemical inhibitors of protein function can be informed by this style of analysis. Our discoveries may hold significant implications for advancing the development of novel treatments for targeting disease mechanisms; they underscore the criticality of investigating the evolutionary past of genes associated with intricate neurological processes. However, further research is needed to fully understand the role of *Rem2* in mammalian biology and evolution.

## Conflicts of interest

The authors declare that they have no competing financial interests that could have influenced the work in this paper.

## Acknowledgements

We thank all our colleagues from the Acme Computational Molecular Evolution Group (ACME) for discussions which helped to improve this paper.

## Supplementary Material

**Figure S1.**
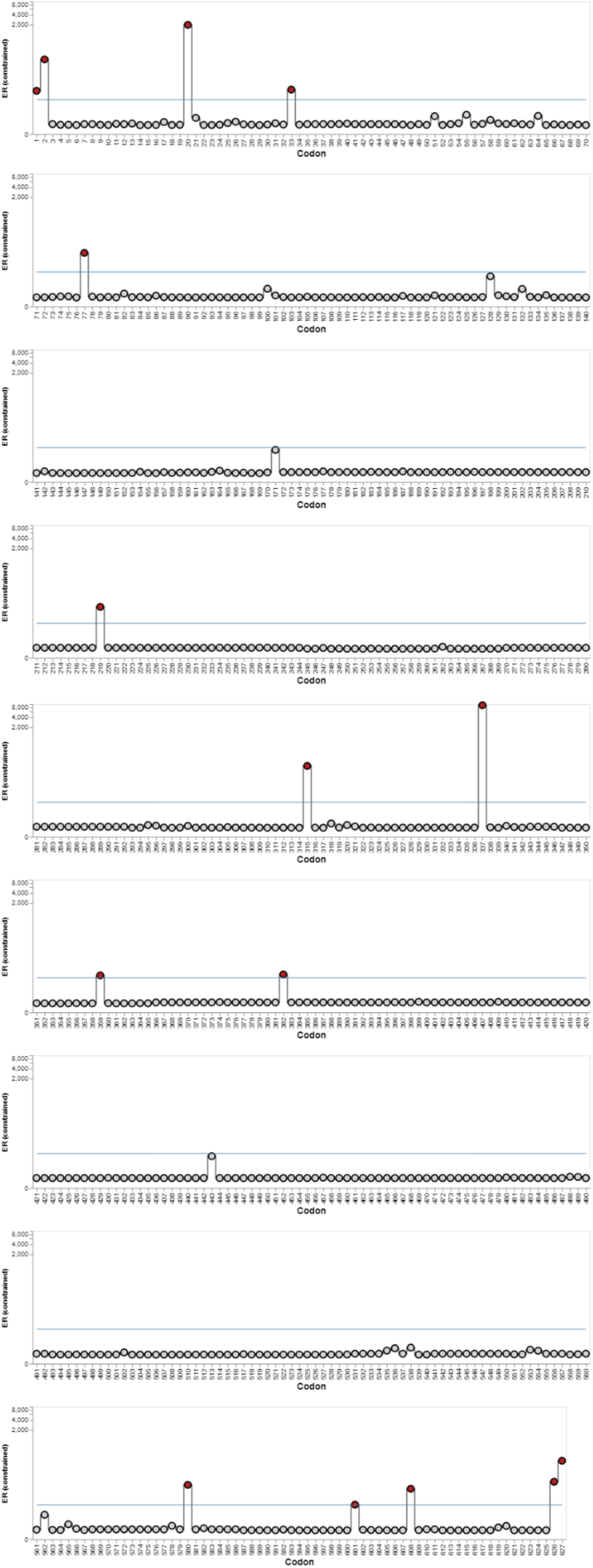
BUSTED+S Evidence ratios (ER, site level likelihood ratios) for ω>1, comparing the unrestricted model with the model where max(ω) := 1, and all other parameters are kept at their maximum likelihood values. Solid line = user selected significance threshold. Values capped at 10000 for readability. We found 15 Sites with ER≥10 for positive selection.

**Table S1.** List of accessions and species used in this study. https://github.com/aglucaci/AOC-REM2/blob/main/data/mammalian_REM2/Rem2_orthologs.csv

**Table S2.** Detailed site-by-site results from the FEL analysis. https://github.com/aglucaci/AOC-REM2/blob/main/tables/mammalian_REM2_FEL_Results_FDR_adjusted_mapped.csv

**Table S3.** Detailed site-by-site results from the MEME analysis. https://github.com/aglucaci/AOC-REM2/blob/main/tables/mammalian_REM2_MEME_Results_FDR_adjusted_mapped.csv

**Table S4.** Detailed site-by-site results from the BGM analysis.https://github.com/aglucaci/AOC-Rem2/blob/main/results/mammalian_Rem2/mammalian_Rem2_BGM_Results.csv

**Table S5.** TM-scores and RMSD calculations from our extinct and extant Rem2 molecular sequences, and TN93 genetic distance calculations. https://github.com/aglucaci/AMES-REM2/blob/main/tables/TM_Align_Results_with_GeneticDistance.csv

## Notes

### Competing Interest Statement

The authors have declared no competing interest.

